# MRI-to-Synthetic 3D Gel Brain: Proof-of-Concept Fabrication For Intra-Parenchymal Diffusion Studies

**DOI:** 10.1101/2022.10.27.514134

**Authors:** Lawrence Ray, Caleb Shaw, Stephanie Otto, Cecile Riviere-Cazaux, Lars Strother, Terry Burns, M. Rashed Khan

## Abstract

Bioprinting technologies utilize hydrogel-based biomaterials to more accurately depict *in vivo* physical conditions within *in vitro* studies, yet, manufacturing the human brain from soft, poroelastic hydrogels remains a fundamental challenge. Conventional manufacturing routes to fabricate hydrogel brain models using techniques, i.e., 3D printing, seems challenging. This study aims to demonstrate an inverse replica molding fabrication technique that can overcome these challenges while maintaining the complex shape of an individual subject’s brain in a miniaturized model—allowing for a more robust hydrogel model that can capture the interactions between diffusing molecules and brain boundaries. This is done by taking a subject’s magnetic resonance imaging (MRI) scan and reconstructing the outer pial surface into a mesh surface. The mesh was then converted to an STL and printed out using an extrusion printer. A silicon mold was made from this print into which agarose was gelled. Once fully gelated, the synthetic gel brain was then carefully removed. Two infusion trials were run in the gel brain, each using a different infusion site. Then a diffusion profile was established and compared to a simple gel infusion model. The result shows different diffusion profiles at each location and between the simple and complex models. This model can better represent the interference the complex shape of the brain has on particle movement compared to simple gel models.

## Introduction

Studies investigating brain tissue diffusion of therapeutics and biomarkers are limited by the central nervous system’s relative inaccessibility and methods for sampling and experimenting within the *in situ* human brains. This project aims to meet the strong clinical need for a translational *in vitro* model capable of accurately modeling the brain’s complex structure for molecular diffusion studies. Recent advances in material sciences and tissue engineering have harnessed synthetic models to empower the simulation, design, repair, and replacement of damaged organs or tissues^1–5^. Prior work has focused on establishing a fundamental understanding, including 3D printing soft materials for making scaffolds of synthetic tissues and organs ^2,6–10^. Despite these advances, a paucity of soft tissue models can be leveraged for diffusion brain modeling studies. Modeling of the human brain is further complicated by the complex nature of its gyri and sulci, each impacting how a molecule diffuses through the cytoarchitecture. Early research in brain modeling demonstrated that 0.6% agarose gel is a prime candidate for modeling diffusion within the brain due to its ability to mimic the consistency of brain parenchyma ^1,11–13^. However, these studies were limited to simple geometric shapes that could not replicate the human brain’s complex structure. Limitations in 3D printing using the brittle material result in the compromised structural integrity of the soft materials. As such, no study has been able to reproduce the complex architecture of the human brain in a synthetic diffusion model until now.

This project innovates a fabrication technique wherein the structure of a patient’s brain can be synthetically modeled by utilizing inverse replica molding from patient-specific MRI data. This approach empowers a low-cost setup to test drugs and molecular diffusion models, which researchers and clinicians can leverage to investigate how the brain’s complex architecture interacts with infused particles. Our study complements the already extensive anatomical models used to improve surgical planning, training, and general education in disease research, treatment planning, cosmetics, orthodontics, neurosurgery, and cardiology^7,8,12,14–17^.

In this project, agarose-based miniaturized synthetic gel brain models were established to mimic the healthy brain environment for a proof-of-concept molecular diffusion study. We acquired the MRI data from the Dallas Lifespan Brain Study, an open repository. Next, we used the automated neuro-analytic/reconstruction software, FreeSurfer, to begin our fabrication process. Once the software reconstructed the brain’s pial surface, a mesh outline of the gray matter was extrapolated and converted to a usable raw STL file format. Then, the raw brain STL was corrected for mesh or volumetric errors in MeshLab. The mesh was then 3D printed into a rigid brain. Ecoflex elastomer was then used to create a mold of the print. Once the print was removed from the mold, 0.6% agarose was poured into the mold to make the synthetic gel brain. The gel brains were then prepped for the fundamental molecular diffusion experiments (Figure 1). Then a simple petri-dish model was made of 0.6% agarose, and the diffusion profiles were compared. The anticipated asymptotic decay in concentration from a single source was found to be perturbed by the presence of the gyroids (folded structure). This further shows that the model’s retention of the brain’s complex structure has a perceived effect on the molecules’ movement through the system, demonstrating that this new synthetic gel brain can better represent human brain boundary interactions in a cost-effective synthetic model.

**Figure 1:**
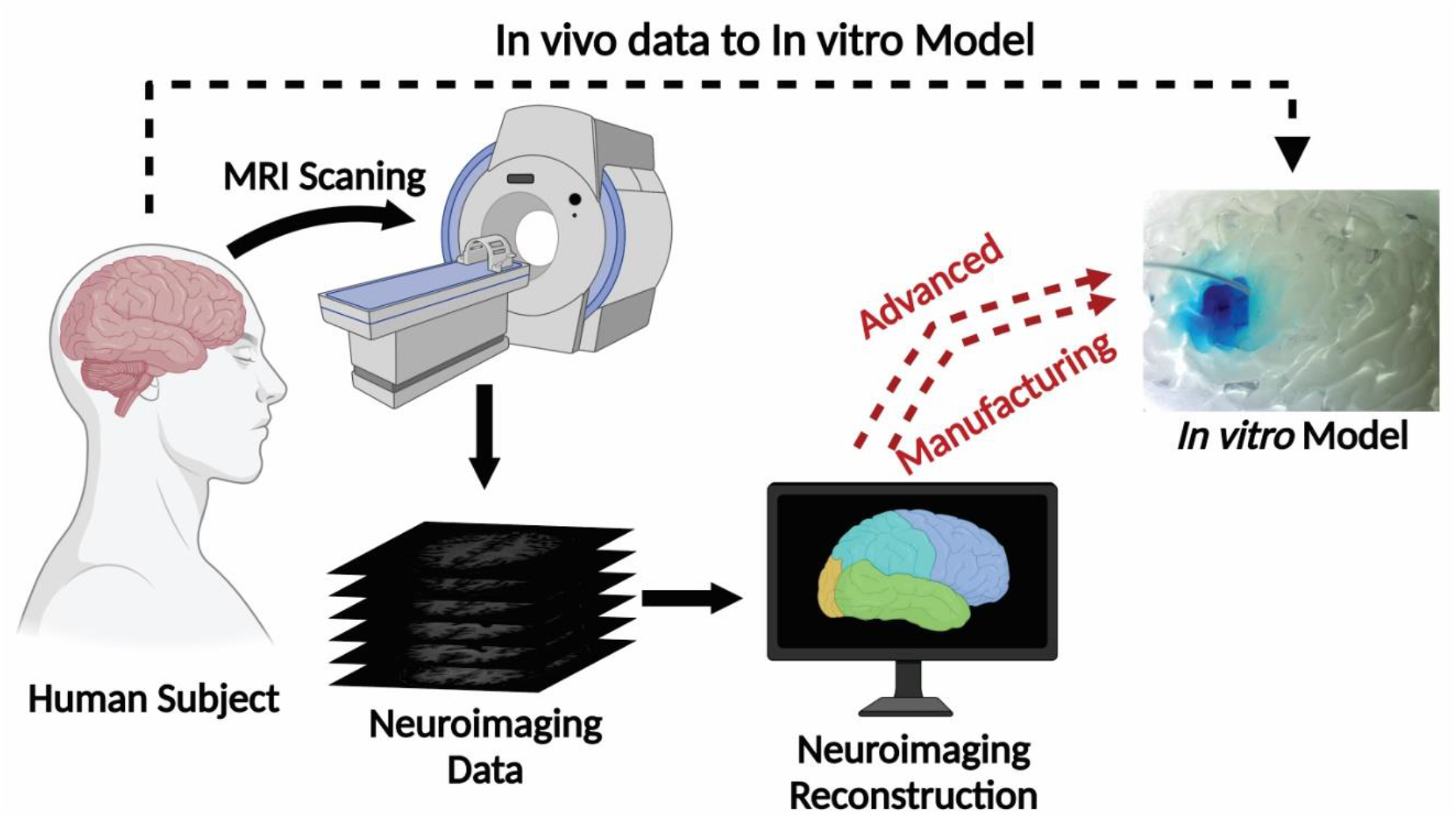
Schematic showing the key steps form collecting patient data to developing an *in vitro* model based off patient data. A healthy brain scan was analyzed and reconstructed into a 3D mesh structure of the pial surface. Advanced manufacturing methods were used to develop a synthetic gel model for *in vitro* diffusion experiments.

## Materials and Methods

### MRI data

MRI data were obtained from the NeuroImaging Tools & Resources Collaboratory site using the Dallas Lifespan Brain Study database, XNAT (nitrc.org). A T1 weighted scan was utilized from a 30-year-old male, with imaging on a 3T Siemens ECAT HR MRI scanner utilizing MPAGE (magnetization-prepared rapid gradient-echo), illustrated in Figure 3A.

### STL extraction

FreeSurfer version 6.0.0 (Stanford University, https://surfer.nmr.mgh.harvard.edu/) was used to analyze and reconstruct the NIfTI-1 (neuroimaging informatics technology initiative) MRI file. The software reconstructed the pial surface, the surface between the cortex and the cerebral spinal fluid, and stored this surface data as a 3D file. This file was then converted to an STL format (using the conversion command shown in Supplementary Figure 2).

### Processing of the STL

MeshLab was used to correct artifacts in the raw STL (https://www.meshlab.net/#description). Quadric Edge Collapse Decimation filter and the Laplacian Smooth tools were used to filter artifacts (Supplementary Figure 1). These filters were applied to these specific locations and the entire mesh. The filters were applied with a setting value of 1 to 2.

### 3D print

A Stratasys uPrint SE Plus 3D printer (Stratasys Inc.) was used to print the reconstructed brain model using acrylonitrile butadiene styrene (ABS) thermoplastics. Printing was performed at DeLeMare Library, University of Nevada, Reno. Default factory parameters with a layer resolution of 0.254mm and minimum wall thickness of 0.914mm were used. The uPrint software was used to reduce the mesh to 40% of its initial size, and the high-density infill option was selected for printing. Each hemisphere took ∼10 hours to print and 4 hours in a lye bath to degrade the water-soluble support structure, made with polyvinyl acetate (PVA) material. The PVA support structure temporarily supports the primary structure. The printing process took ∼20 hours to get the final structure of the whole brain, followed by 8 hours of cleaning in the lye bath. The approximate material cost per hemisphere was $14, with two individual hemispheres and one whole brain costing approximately $28. The scheme of this low-cost advanced manufacturing process is shown in Figure 2. The thermoplastic physical prints are shown in Figure 3B.

**Figure 2:**
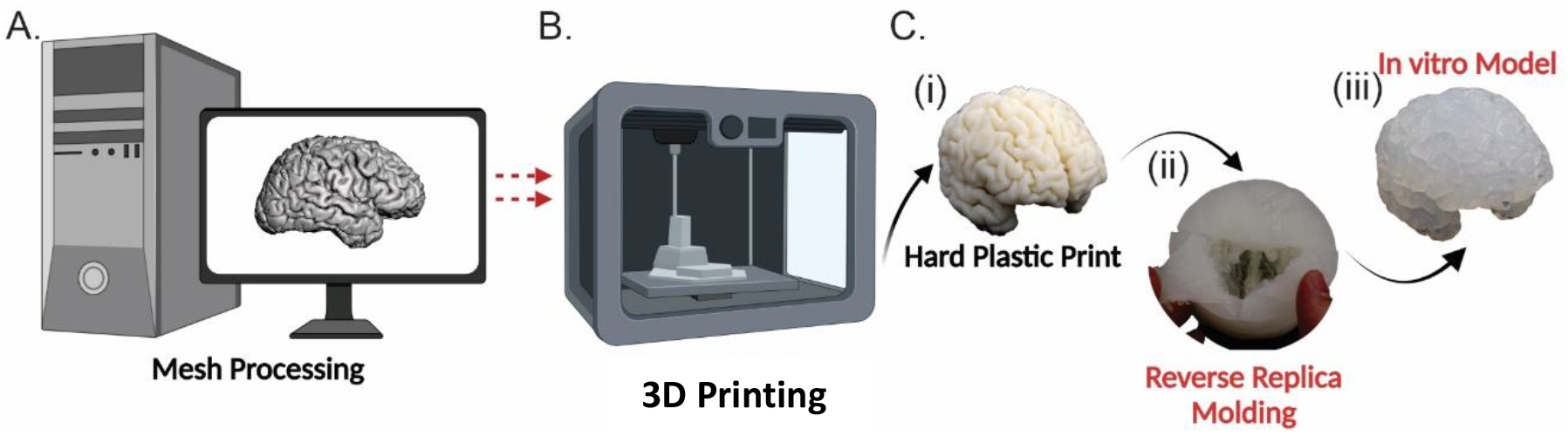
Schematic showing the advanced manufacturing methods. A. Mesh processing of the reconstructed brain. B. Using a thermoplastic printer, a physical model of the brain. C. Prints were created as follows: (i) ABS hard plastic brain print, (ii) inverse replica molding using silicone on the hard plastic print, and (iii) the in vitro model of the brain was created from 0.6% agarose.

**Figure 3:**
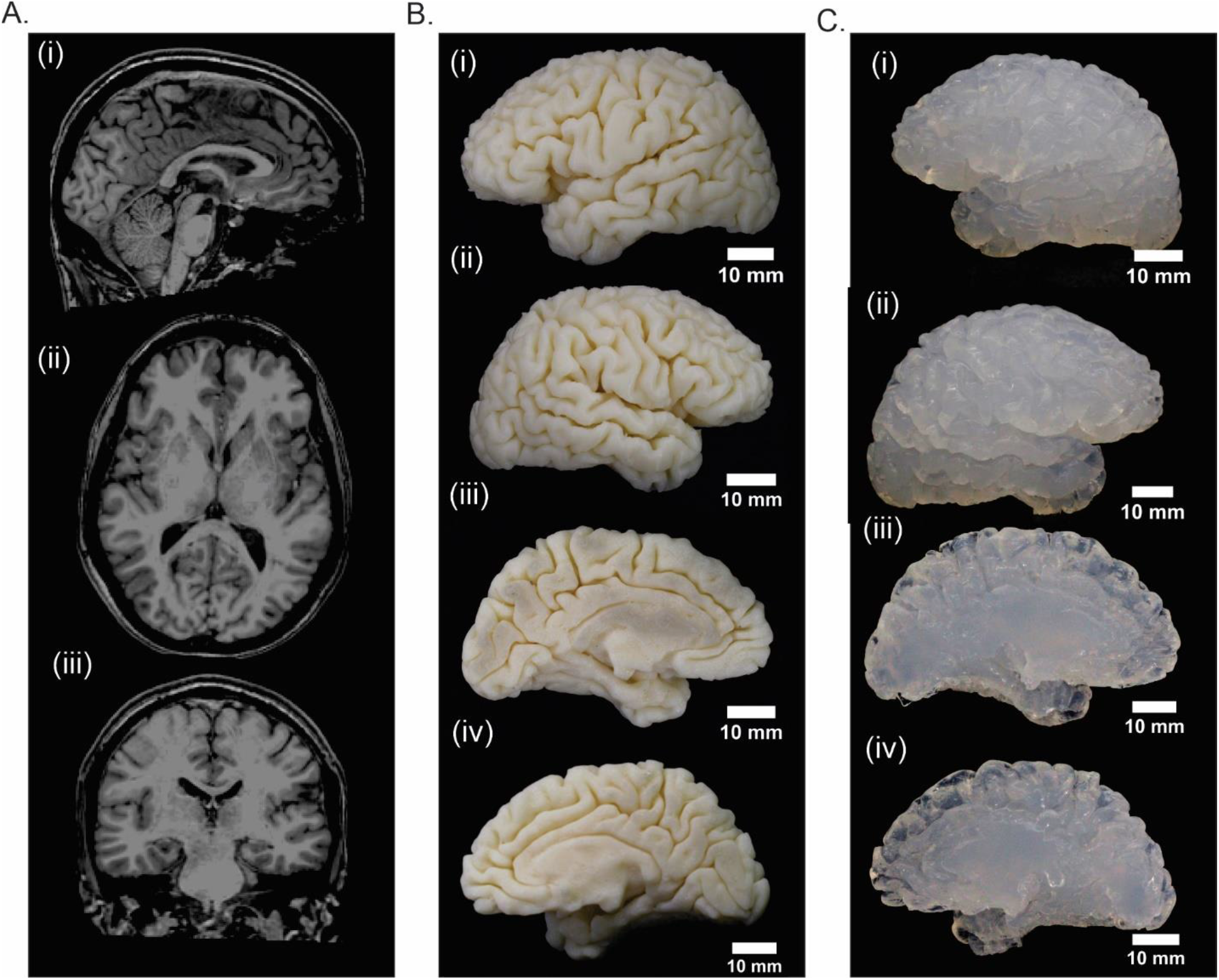
Side by side comparison of original MRI scans, 3D printed brains, and synthetic brains. A. (i)Sagittal, (ii)Axial, and (iii)Coronal slices of the original MRI scans. B. From top-down, the 3D printed brain is shown at different viewpoints starting with the (i) lateral side of the left hemisphere, (ii) lateral side of the right hemisphere, (iii) medial side of the left hemisphere, and (vi) medial side of the right hemisphere. C. From top-down, the soft brain is shown at different viewpoints starting with the (i) lateral side of the left hemisphere, (ii) lateral side of the right hemisphere, (iii) medial side of the left hemisphere, and (vi) medial side of the right hemisphere.

### Casting

The 3D-printed brain was then coated with a non-sticking agent (corn oil) before being placed in EcoFlex PLATINUM CURE SILICONE RUBBER (SCS Direct Inc.) to cast the inverse mold. The mold was then set in a non-stick container at room temperature. The mold was checked every 30 minutes to verify if it was set properly. After approximately 6 hours, the full brain print was separated from the silicone coating by making a single cut across the right hemisphere, from the frontal lobe to the occipital lobe. The single-hemisphere molds were cut from the frontal lobe to the occipital lobe, following the axial plane inferior to the corpus callosum. Each mold was submerged in the non-sticking agent and sonicated for 5 minutes. The silicone was then carefully peeled off the printed brain to avoid damage to the more delicate brain sulci. Once fully removed, the mold was cleaned with soap and water to remove any non-sticking agent from the mold, following a final rinse with deionized water. Three molds were manufactured: left hemisphere mold, right hemisphere mold, and full brain mold (Figure 2B).

### Synthetic Brain Fabrication

The synthetic brains were created using 0.6% agarose, which mimics the poroelastic properties of the normal mammalian brain. A 50-milliliter solution of 0.6% agarose was mixed and heated, using a VWR Hotplate and Stirrer, at 140°C at 160 RPM until the solution was clear. It was then degassed by an AR-100 Conditioning Mixer by Thinky Inc. for 25 seconds. Then the agarose was poured into the whole brain mold; the mold was left to polymerize for approximately 45 minutes at 4°C. The single-hemisphere molds were left to solidify at 4°C for approximately 30 minutes. The gel brains were removed from the mold and left at room temperature, for approximately 20 minutes, before infusion.

### Simple Petri Dish Model

The simple gel brain model was created in a small petri dish using an agarose concentration of 0.6%. A 25ml solution of 0.6% agarose was mixed and heated, using VWR Hotplate and stirrer, at 140°C at 160 rotations per minute until the solution was clear. It was then degassed by an AR-100 Conditioning Mixer by Thinky Inc. for 25 seconds. Then poured into a small petri dish and left to gelate.

### Comparing Size

MeshLab was used to measure the select brain regions, and then the measurements were reduced to 40% of their initial measurements. Then the gel brain was measured using calipers in the select regions measured in MeshLab. These measurements were recorded in supplementary Table 1.

### Molecular Diffusion Study

An Olympus SZ61 stereoscope with a 10-megapixel camera was used to measure particle movement within the synthetic gel brain’s poroelastic matrix over time using methylene blue (Quality Laboratory Solutions and Chemicals, Reno), with a molecular weight of 319.85 g/mol. For infusion, an open-ended high-density polyethylene (HDPE) tube was used. Its diameter was 0.28 mm. The first two trials were in the synthetic gel brain using two insertion sites with a probe depth of approximately 5 millimeters. The first probe insertion site was in the middle frontal gyrus inferior segment of the synthetic brain and the second insertion site was in the superior frontal gyrus superior segment. The second trails gel brain was sliced, bisecting the probe’s insertion site, producing two sagittal slices, lateral and medial, with an approximate thickness of 5 mm. Then the petri dish gel model was used for a similar infusion study, using the same probe at a depth of approximately 5mm. The probe was inserted through the bottom of the petri dish. The duration of the infusions was 10 hours, and we used a NanoJet pump (Chemyx Inc.) at an infusion rate of 18 µl per hour. The infused solution had a methylene blue concentration of 1.25 mol/m^3^. During the infusion, images were taken every minute for 600 minutes. The time points of 150-minutes, 300-minutes, 450-minutes, and 600-minutes during all infusions were analyzed using surface analysis line scans, and the intensities were graphed (Figure 4 and Supplementary Figure 3).

**Figure 4:**
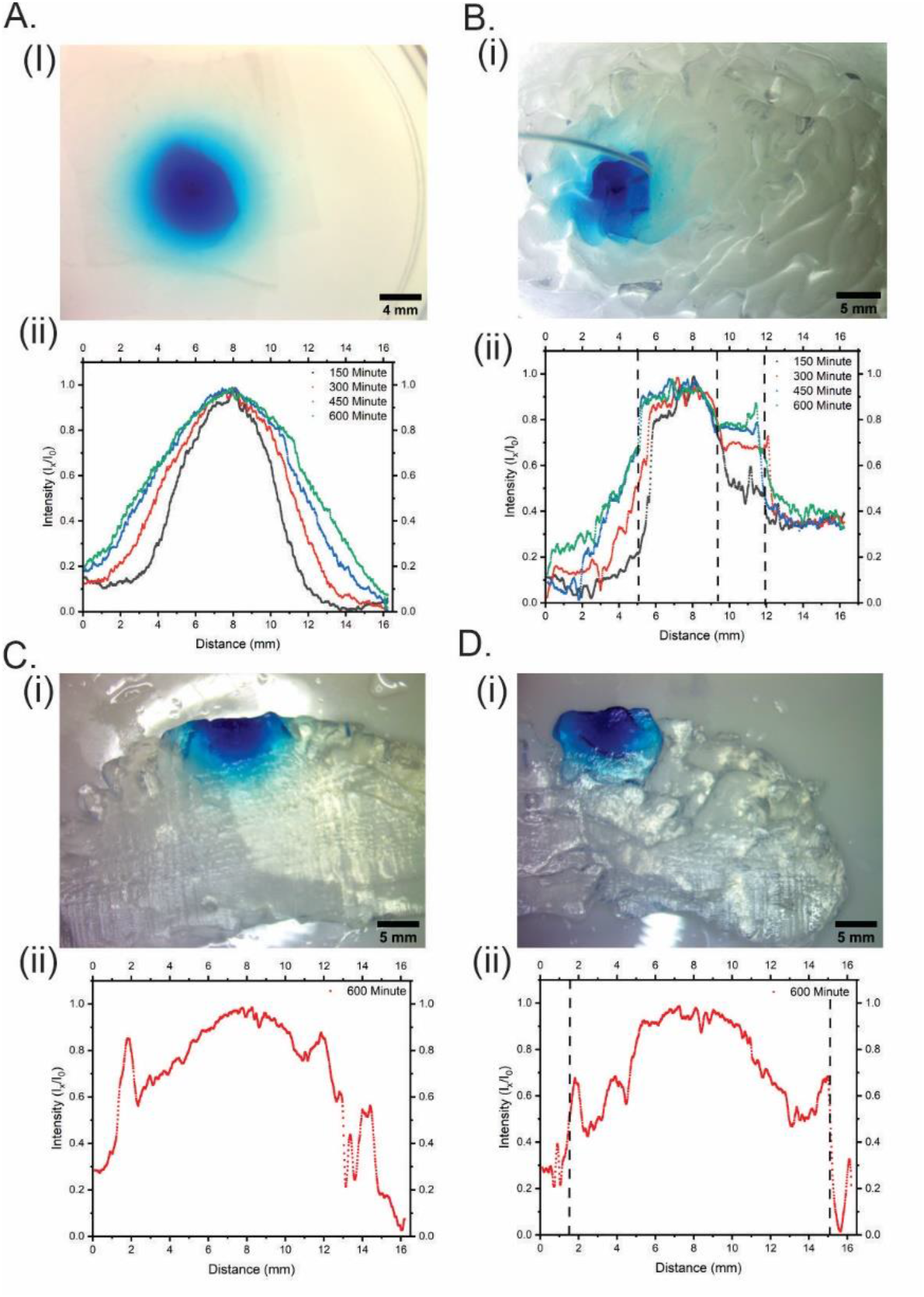
Surface analysis of methylene blue’s intensity in the synthetic brain and petri-dish models. A. (i)Surface analysis of the petri dish at time point 600-minute, (ii) surface plot of the petri dish at time points 150-minute, 300-minute, 450-minute, and 600-minute. B. (i) Soft brain infused in the middle frontal gyrus, (ii) surface line plot of the infusion at the middle frontal gyrus at time points 150-minute, 300-minute, 450-minute, and 600-minute with sulci depicted by the dashed line. C. (i) Sagittal slice bisecting infusion site on the lateral side, (ii) surface plot of the infusion site at time point 600-minute. D. (i) Sagittal slice bisecting infusion site of the medial side, (ii) surface plot of the infusion site with sulci represented by the dashed lines.

## Result

### Experimental Paradigm

A patient-specific synthetic brain phantom was fabricated using agarose gel (Figure 2B). This model was then infused with a drug mimic, methylene blue, and its diffusion pattern was then analyzed.

### Comparison of Different Infusing Regions

Both infusions into the synthetic gel brains were recorded and analyzed. The relative intensity of both graphs (Figure 4A. and Figure 4C.) had distortions in their diffusion profiles compared to each other. This distortion was observed at the sulci and gyri depicted in the surface plot profiles (Figure 4A (ii) and Figure 4C (ii)).

### Comparison of the Synthetic Gel Brain and Petri Dish Models

All infusions used methylene blue at a concentration of 1.25 mol/m^3. The probe insertion site is located in all plots at the 8-millimeter distance (Figure 4 and Supplementary Figure 3). The infusions into the synthetic gel brains were at a rate of 18 µL per hour in the middle frontal gyrus inferior segment and superior frontal gyrus posterior segment. The probe depth was 5 mm, and the infusions lasted 10 hours (180 µL total volume). The 150-minute, 300-minute, 450-minute, and 600-minute images were then analyzed from the model’s surface, and a relative diffusion gradient was established after the moving averages were taken. In the middle frontal gyrus infusion, the relative intensity of methylene blue in the infusing gyri was measured at ∼0.9, between a distance of ∼5-9 mm (Figure 4B (ii)). The sulci were then observed interacting with the methylene blue diffusion at distances ∼5 mm, ∼9 mm, and 12mm. These interactions decrease the relative intensity by more than 0.5 in the first millimeter from the sulcus at a distance of ∼5mm in Figure 4B (ii). The sulcus at a distance of ∼9mm recorded a relative intensity drop of ∼0.1 to ∼0.4 between distances ∼9mm to ∼12mm. The last sulcus was seen between distances 12mm and 16mm decreasing the intensity to approximately 0.4. For time points 300-minute, 450-minute, and 600-minute.

The second synthetic gel brain relative intensities were shown to be at the highest in the infusing gyrus, between distances ∼4mm to ∼13mm. The relative intensities reported were in the range of ∼1.0 to 0.6. The intensity dropped by ∼0.1 after sulci at the 13 mm distance (Supplementary Figure 4 (ii)). Little change was seen after the sulci at the ∼4 mm distance.

The synthetic gel brain slices (Figures 4C and D) were cut at a thickness of ∼5 mm on the sagittal plane. The slices were then imaged, and a diffusion profile was taken from a line scan perpendicular to the probe insertion site. The diffusion profile of the lateral slice (Figure 4C (ii)) had no distinct sulci when the line scan was performed on the slice image (Figure 4C (i)). The medial slice diffusion profile (Figure D (ii)) is seen to have interfering sulci at distances of ∼1.5 mm and ∼15 mm. At these sulci, the intensities spiked, followed by a decreasing intensity range from ∼0.4 to ∼0.7.

The petri dish infusion was analyzed at time points 150-minute, 300-minute, 450-minute, and 600-minute, and a diffusion profile was established. The diffusion curve peaks at ∼8mm, where the probe was inserted, and an asymptotic decay is observed (Figure 4A (ii)). The difference between the diffusion profiles in Figure 4 was due to structural variation between models. The synthetic gel brain shows interference from the complex structure compared to the normal diffusion curve in the petri dish.

### Size Comparison of Mesh Model to Synthetic Gel Brain

The mesh size was compared in 11 locations, after being reduced to 40% of its initial size, to the same locations in the synthetic gel brain. The average difference in the size of all locations was 0.6 mm.

## Discussion and Conclusion

In this study, a low-cost and efficient agarose-based, MRI-derived synthetic gel brain was designed, fabricated, and investigated to address the unmet need for a translationally relevant *in vitro* model for molecular diffusion studies. The subject’s brain was recreated from an MRI scan using replica molding on a 3D-printed model. This novel approach enabled a soft brain reconstruction of the complex shape of the human brain in 0.6% agarose gel material to most accurately mimic normal brain parenchyma ^1^. With this miniaturized model in hand, future studies will investigate how a complex shape, like the brain, impacts particle and drug movement and how such data may be leveraged to understand better and interpret biomarker and candidate therapeutic drug levels in regions of interest (ROI).

As a proof-of-principle study, surface analysis of the soft brain model after a 10-hour infusion revealed the differential movement and interaction of methylene blue with the brain’s gyri and sulci compared to the normal distribution curve of 0.6% agarose (Figure 4). The surface analysis revealed that methylene blue’s movement through the material was impeded by the presence of sulci separating each gyrus. As the line plot moved from one gyrus to another, separated by a sulcus, methylene blue intensity decreased, suggesting an impediment of movement by these anatomical structures. This demonstrates that interfering sulci impact the diffusion of particles throughout the system. This *in vitro* approach empowers a previously unprecedented opportunity to evaluate diffusion in an individualized synthetic agarose model that closely resembles brain parenchyma.

A shortcoming of this study is the experimental measurement technique, which applies a 2D line scan onto a 3D object, limiting the full quantification of dimensional movements across the model as would occur *in vivo*. Improved measurement techniques may better portray the experimental results to understand how complex anatomical structures impact a dimensional phenomenon such as diffusion within the human brain. Ongoing studies using a particle that can be traced in MRI while the infusion is actively taking place would enable the 3D extraction and quantification of real-time 3D diffusion data in a clinically relevant paradigm. We envision this novel approach will be of great use for future studies examining biomarkers and candidate therapeutics collection and interpretation within a feasible and individualized *in vitro* model.

## Acknowledgment

MRK conceived the idea. MRK and TB supervised the project. MRK and LS consulted and extracted neuroimaging support through Integrative Neuroscience at UNR. SO trained LR to run FreeSurfer analyses. CS experimented with LR. LR executed and ran the experiments. MRK, TB, CRC, LR, and CS edited the drafts. MRK acknowledges the funding support of the VPRI startup fund. We also acknowledge the contributions of Braden Norris, Gerra Licup, Dylan Obrochta, Jon, Jay Bemel, PJ Hall, and Trevor Haub from the University of Nevada, Reno, for initially collecting experimental data in agarose diffusion experiments. CRC contributed to the writing. CS aided in analyzing results. LR designed, executed, and experimented with the project, analyzed the results, and wrote the manuscript. Facilities were accessed through Integrative Neuroscience Department for computation and neuro-analysis/reconstruction of the data through the support and supervision from LS and affiliation of MRK. Neuro-analysis and reconstruction guided by SO. The data was acquired from NeuroImaging Tools & Resources Collaboratory (NITRC) Dallas Lifetime Study subjects, funded by NIH grant numbers 2R44NS074540 and 1U24EB023398.

